# Phylogenetic analysis of West Nile Virus in Maricopa County, Arizona: Evidence for dynamic behavior of strains in two major lineages in the American Southwest

**DOI:** 10.1101/223503

**Authors:** Crystal M. Hepp, Jill Hager Cocking, Michael Valentine, Steven J. Young, Dan Damien, Krystal Sheridan, Viacheslav Y. Fofanov, Joseph D. Busch, Daryn E. Erickson, Ryan C. Lancione, Kirk Smith, James Will, John Townsend, Paul S. Keim, David M. Engelthaler

## Abstract

West Nile Virus (WNV) has been detected annually in Maricopa County, Arizona, since 2003. With this in mind, we sought to determine if contemporary strains are established within the county or are annually imported. As part of this effort, we developed a new protocol for tiled amplicon sequencing of WNV to efficiently attain greater than 99% coverage of 14 WNV genomes collected directly from positive mosquito pools distributed throughout Maricopa County between 2014 and 2017. Bayesian phylogenetic analyses revealed that the contemporary genomes fall within two major lineages, NA/WN02 and SW/WN03. We found that all of the Arizona strains possessed a mutation known to be under positive selection (NS5-K314R), which has arisen independently four times. The SW/WN03 strains exhibited transient behavior, with at least 10 separate introductions into Arizona when considering both historical and contemporary strains. However, NA/WN02 strains are geographically differentiated and appear to be established in Arizona, with likely origins in New York. The clade of New York and Arizona strains looks to be the most ancestral extant lineage of WNV still circulating in the United States. The establishment of WNV strains in Maricopa County provides the first evidence of local overwintering by a WNV strain over the course of several years in Arizona.

## Introduction

The first human cases of West Nile Virus (WNV) were identified in New York City during the summer of 1999. This mosquito-borne virus belongs to the Japanese encephalitis complex (genus *Flavivirus*) and became well established throughout the United States by 2004. Over a decade later, WNV is still the most important arbovirus nationwide, causing 95% of arboviral diseases reported to the Centers for Disease Control and Prevention (CDC) [1]. The remaining cases are caused by: La Crosse virus, St. Louis encephalitis virus, Jamestown Canyon virus, Powassan virus, eastern equine encephalitis virus, unspecific California serogroup virus, and Cache Valley virus [1]. From 1999-2015 there have been approximately 43,937 human cases and 1,911 deaths associated with WNV infections reported to the CDC [2]. Of those cases, nearly 18% (n=7,764) have occurred in the southwestern states of Nevada, California, Utah, and Arizona [1]. Maricopa County, the most populous county in Arizona, and the 4^th^ most populous county in the United States, detected its first WNV-positive bird (*Passer domesticus* – House sparrow) in September 2003. A positive mosquito pool was detected one week later, followed shortly thereafter by the first autochthonous human case in November of that same year. Human WNV cases within Maricopa County peaked dramatically in 2004 with 355 human cases and 14 deaths [3]. A lesser spike in human infections occurred in 2010, with 155 reported cases. While WNV is not presently affecting the human population of Maricopa County to the extent it did in 2004 and 2010, the virus has reliably infected humans in the area each year since its first detection. As of November 17^th^, 2017, there were already 91 confirmed or probable cases in the county, with 89 of those being neuroinvasive cases, and 220 positive mosquito pools (74% were *Culex quinquefasciatus* pools) of 10,228 tested across the Phoenix Metropolitan area [4] (Fig. 1). Given the annual resurgence of WNV in Maricopa County, the purpose of the study presented here was to answer the following overarching question: Are Maricopa County WNV populations established residents or are they reintroduced from other foci annually?

**Figure 1:**
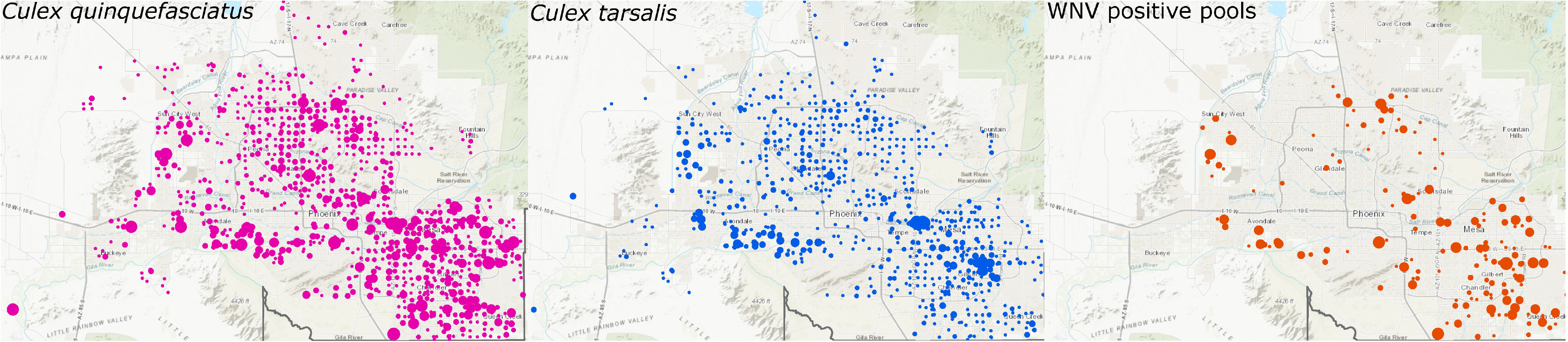
Distribution of WNV and its vectors across Maricopa County in 2017. The dots represent the count of individual female *Culex quinquefasciatus* (left), *Culex tarsalis* (center), or WNV positive mosquito pools (right) at each of 787 carbon dioxide traps distributed primarily throughout the urban portion of the county. Larger circles indicate a higher density at a particular trap.

## Materials and Methods

### Sample Collection

The Maricopa County Environmental Services Department Vector Control Division conducts year-round mosquito surveillance and abatement activities throughout Maricopa County. Mosquitoes are collected once weekly from 787 routine carbon dioxide trap locations distributed throughout the Phoenix metropolitan area (Fig. 1); collections were subsequently sorted by species and sex. Up to 5 pools of 50 individual female mosquitos are pooled and tested for WNV once weekly, using the protocol described in Lanciotti et al. [5]. We selected 14 WNV positive mosquito pools, distributed geographically (Fig. 2) and temporally (from 2014 to 2017, Table S1), for whole genome tiled amplicon sequencing using a novel protocol, based on the method of Quick et al. [6] developed for Zika virus sequencing.

**Figure 2:**
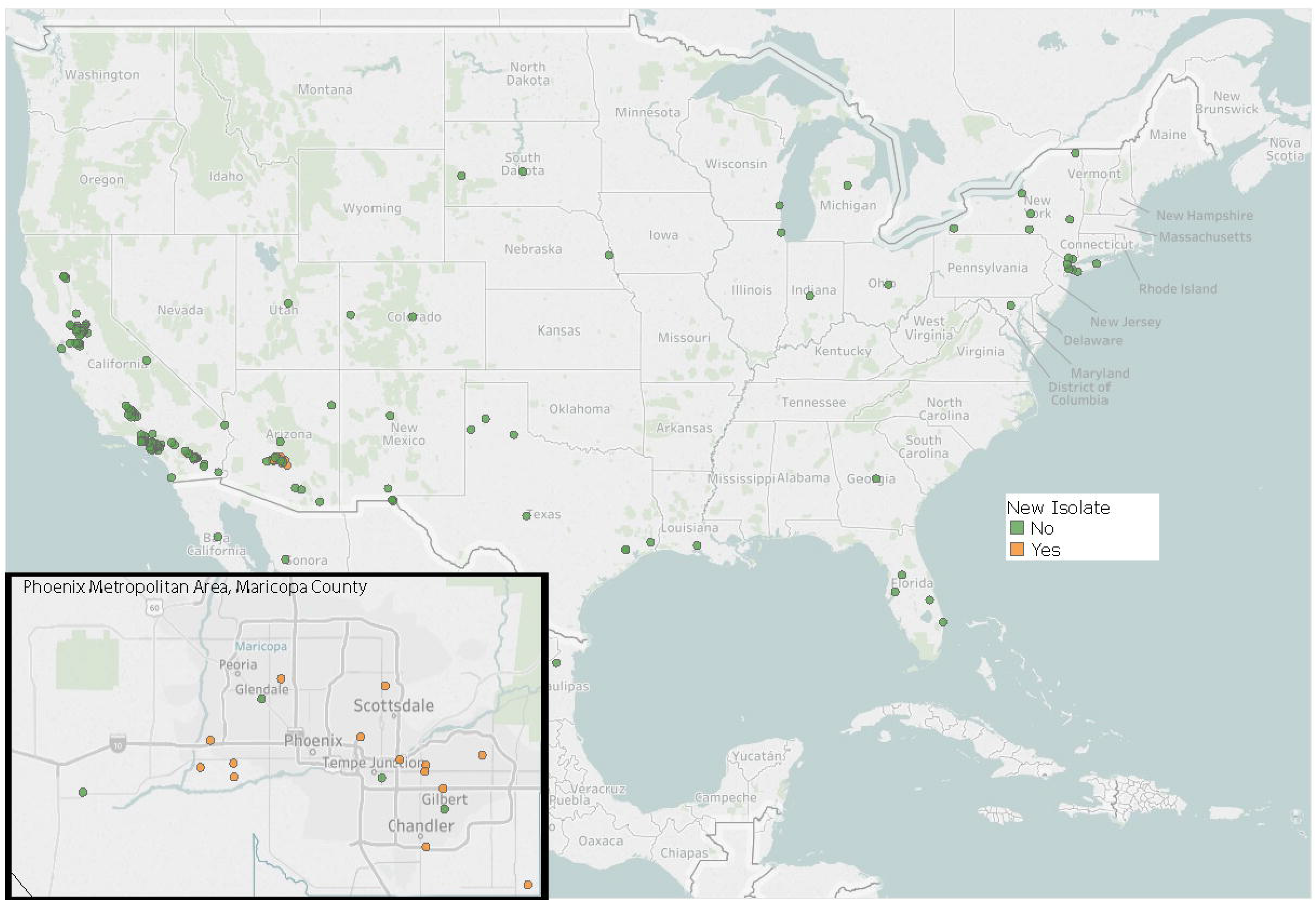
Geographic distribution of WNV strains included in the phylogenetic analysis, except for the New York strains published in Ehrbar et al., [16] where latitude and longitude coordinates were not available. Green points indicate publicly available strains while orange points indicate strains that were sequenced as part of this study. Additional metadata can be found in S1 Table.

### Sample Processing, a new WNV Tiled Amplicon Protocol, and Next Generation Sequencing

Prior to transporting the samples, an equal part of DNA/RNA Shield™ 2X Concentrate (Zymo Research) was added to each mosquito pool to stabilize the RNA and inactivate WNV. The pools consisted of mosquitos in approximately 1.3 ml of TE buffer (Invitrogen, AM9858). All mosquito pools were stored at −80 degrees Celsius prior to extractions. Both DNA and RNA were extracted from the pools in a Biosafety Cabinet using the Quick DNA/RNA Pathogen Miniprep™ kit (Zymo Research). RNA was reverse transcribed into cDNA using the protocol as described in Quick et al. [6], and the product was stored at - 20 degrees Celsius.

The Primal Scheme primer designer software [6] was used to design the multiplex PCR primers for our tiled amplicon sequencing protocol. A total of 41 primer pairs with attached universal tails, split into two pools, were used to amplify regions averaging 372 bases in length, covering the NY99 (NC_009942.1) genome positions 8-10877 (Table S2). The preparation of the tiled amplicons was carried out as described by Colman et al. [7]. For the initial amplification of specific regions of the West Nile Virus genome, each sample was prepared with two pools of primers. Each of the two PCRs per sample consisted of 12.5 μl of KAPA 2G Fast Multiplex PCR Mastermix (Kapa Biosystems, Wilmington, MA), primers from pool 1 or 2 for a final concentration of 0.2 μM each primer, and 2.5 μl of cDNA. The PCR was performed as follows: 3 minutes of denaturation at 95°C, 30 cycles of 95°C for 15 seconds, 60°C for 30 seconds, 72°C for 1 minutes, and a final extension of 72°C for 1 minute. The PCR products were cleaned using 1X Agencourt AMPure XP beads (Beckman Coulter, Indianapolis, IN). Illumina’s sample-specific index and sequencing adapters were added during a second PCR utilizing the universal tail-specific primers. This reaction was prepared with 12.5 μl of 2X Kapa HiFi HotStart Ready Mix (Kapa Biosystems), 400 nM of each forward and reverse indexed primer, and 2 or 4 μl the cleaned and amplified WNV product. This PCR was performed as follows: 98°C for 2 minutes, 6 cycles of 98°C for 30 seconds, 60°C for 20 seconds, 72°C for 30 seconds, and finally 72°C for 5 minutes. The indexed PCR products were again cleaned with 1X Agencourt AMPure XP beads (Beckman Coulter). The amplicon libraries from each WNV sample were quantified using the Kapa Library Quantification kit (Kapa Biosystems) and pooled in equal concentrations. Sequencing was carried out on the Illumina Miseq sequencing platform, using a v2 500 cycle kit.

### Sequence Data Processing

Sequencing reads were aligned to the NY99 Lineage 1 WNV genome (NC_009942.1) using samtools [8] and Bowtie2 [9]. Alignments were visualized in the Integrative Genomics Viewer (IGV), and all sequenced positions had at least 100x coverage [10, 11]. Consensus sequences were exported from IGV, where at least 80% of reads at a position had to have the majority allele for a position to be called an A, T, G, or C. For sites where the majority allele was not represented in at least 80% of the reads at that position, sites were coded according to the International Union of Pure and Applied Chemistry (IUPAC) nucleotide codes. The 14 new consensus genomes and associated metadata were deposited into Genbank using Bankit (Accession numbers: MG004528-MG004541).

### Phylogenetic Analysis with an Incorporated Timescale

Using MUSCLE in MEGA7.0 *[12]*, we aligned a total of 246 genomes, including: i) newly sequenced genomes from this study (n=14); ii) United States-based whole genomes that were included in Pybus et al. [13] (n=104); iii) Arizona-based genomes that were published in Plante et al. [14] (n=3); iv) Southern California-based genomes published in Duggal et al. [15] (n=112); and v) New York-based genomes [16] (n=13) (Table S1). Genomes were selected based on availability of sample metadata, including time and geolocation of collections, except for the New York genomes from Ehrbar et al., [16], where geolocation coordinates were not available. To determine if the WNV genomes in those data exhibited a strong molecular clock signal, we constructed a neighbor joining tree in MEGA7.0 [12] and uploaded the newick file, with associated collection dates into TempEst [17]. The coefficient of determination revealed that nearly 90% (R^2^ = 0.8871) of the variation in root to tip distance can be explained by time. This indicated that a molecular clock analysis would be appropriate for the dataset.

We employed a Bayesian molecular clock method implemented in the BEAST v1.8.0 software package to estimate evolutionary rates for WNV, as well as divergence times for Arizona lineages, [18]. Substitution model selection was carried out in MEGA 7.0.9 for the 246 genomes included in our dataset. The corrected Akaikes’s Information Criterion and Bayesian Criterion results indicated that the General Time Reversible model with incorporation of a gamma distribution of among-site rate variation would be the best fitting for the dataset. To determine the best fitting clock and demographic model combinations for these data, path sampling [19] and stepping stone [20–22] sampling marginal likelihood estimators were employed to compare the Uncorrelated Lognormal (UCLN) clock model combined with the Bayesian Skyline model as used in Pybus et al. [13], the Gaussian Markov Random Field Bayesian Skyride model [23], or the Bayesian Skygrid model [24]. Each of the three model combinations were iterated 100,000,000 times, where each Markov chain was sampled every 10,000 generations. We found that the UCLN Bayesian Skyline combination outperformed the other two (Table S3). Using the UCLN Bayesian Skyline model, we ran three additional chains for 100,000,000 generations, sampling every 10,000, and found convergence within and among chains using Tracer v1.6 [25]. We used LogCombiner to merge the four different chains, discarding the first 10% as burn in (40,000,000 generations), and then resampled every 36,000 generations. The resulting file was input to TreeAnnotator to produce a maximum clade credibility tree, and then visualized using Fig Tree v1.4.3 [26].

### Defining WNV lineages

The three major lineages of WNV in the United States have been defined by two nonsynonymous mutations (E-V159A, NS4A-A85T) within the polyprotein, and we have identified the locations of these mutations on the reconstructed phylogeny reported here (S1 Table, Fig. S1). Members of the NY99 lineage, the most ancestral lineage of WNV that entered North America, possess an valine at position 159 of the envelope protein, while nearly all isolates collected after 2002 have an alanine at that position, and are part of both the North American/WN02 (NA/WN02) and distal Southwestern/WN03 (SW/WN03) lineages [27, 28]. The entire SW/WN03 genotype is defined by the NS4A-A85T mutation. The more distal SW/WN03 taxa additionally possess the NS5-K314R mutation [29], which also independently arose in a clade of Southern California strains residing within the NA/WN02 lineage (S1 Table and Fig. S1) [15].

## Results and Discussion

In an effort to better understand the recent dynamics of West Nile Virus circulation in the Phoenix metropolitan area of Maricopa County, Arizona, we sequenced 14 genomes, varying in location within the county (Fig. 2 inset), sampling date (2014-2017), and vector species (S2 Table). We evaluated these strains in a nationwide context, by performing a Bayesian phylogenetic analysis (Fig. 3, Fig. S1). This included comparisons to an additional 232 genomes which were distributed throughout the United States (Fig. 2), in order to estimate: 1) when WNV was first introduced into Maricopa County, AZ, 2) how many distinct WNV introductions have occurred, 3) if contemporary strains belong to lineages that have become established in Maricopa County, and 4) the temporal span of such establishment.

**Figure 3:**
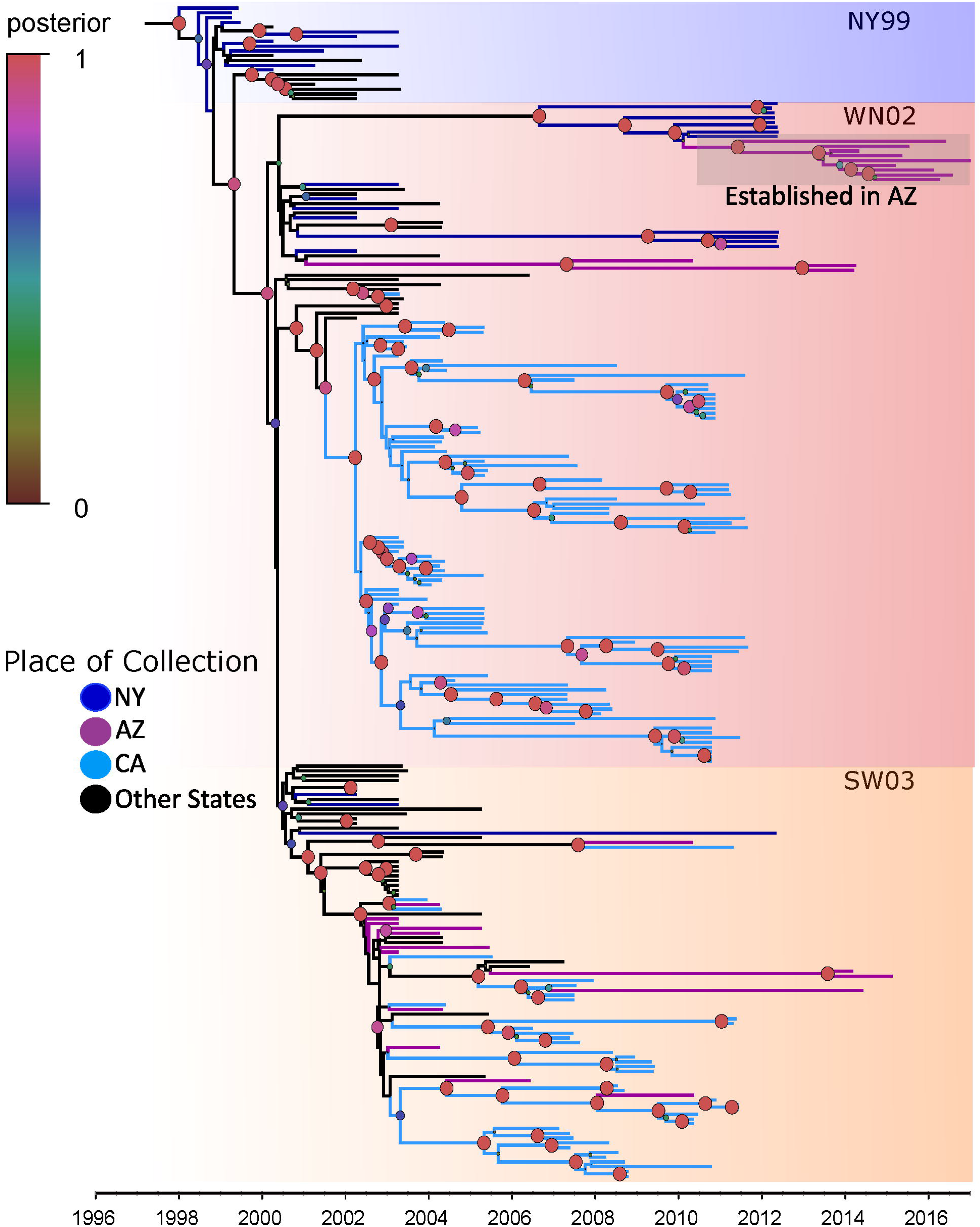
Maximum clade credibility phylogenetic tree reconstructed from 246 nationally distributed WNV genomes. The gradient squares indicate the phylogenetic position of the NY99 (blue), NA/WN02 (red), and SW/WN03 (orange) clades. Posterior probabilities are represented by the size (larger circles have higher values) and color (defined by the color legend) of circles at each node. The established clade in Maricopa County is encompassed by a grey shadow.

The phylogeny revealed that WNV strains in Arizona have been genetically diverse, are represented in both major lineages (NA/WN02 and SW/WN03) that are still circulating in the United States (Fig. 3 and Fig. S1). The placement of Maricopa County strains within the NA/WN02 and SW/WN03 lineages was first described in Plante et al. [14], and in agreement with that study, we find that 11 of the genomes sequenced here belong to the NA/WN02 lineage, while 3 others cluster within the SW/WN03 lineage.

Maricopa County strains interspersed within the SW/WN03 clade exhibit polyphyletic clustering, and are clustered with strains sampled from California, Colorado, Texas, and New Mexico. This clustering behavior indicates that the lineage has dispersed extensively throughout the southwestern United States. Furthermore, the most recent sampling was in 2015, suggesting that this lineage is still in circulation. The strains belonging to the SW/WN03 lineage were the first to be introduced into Arizona, during 2002 (mean estimate: 2002.48, 95% CI: 2002.42-2002.99) (Fig. 3 and Fig. S1), and have been reintroduced on at least nine occasions represented by only one or two strains each time. The contemporary strains sequenced in this study (MG004533, MG004540, and MG004537) represent two introductions that are part of a small clade, which includes strains from Southern California and Texas. This may indicate that the genomic diversity of the SW/WN03 lineage in Arizona has been reduced in recent years.

The NA/WN02 lineage was first detected in Arizona in 2010 [14], and the strain representing the first detection (KF704158.1) clusters monophyletically with two additional contemporary strains that were collected in 2014 (MG004529 and MG004539) (Fig. 3 and Fig. S1). The estimates of time to most recent common ancestor (TMRCA) for those strains indicate that the initial introduction occurred in 2007 (mean estimate: 2007.31, 95%CI: 2005.86-2009.23), three years prior to first detection. While we find that strains from Arizona are dispersed throughout the tree, 9 of the 14 strains sequenced in this study clustered monophyletically, with a most recent common ancestor entering Arizona in 2011 (mean estimate: 2011.42, 95%CI: 2010.63-2012.7). This monophyletic clade, which includes strains collected from 2014 through 2017, is nested within a paraphyletic clade of New York strains that were collected in 2012, and indicates a single introduction into Arizona of the New York lineage. This monophyly and node dating indicates both a single and recent introduction directly from New York to Arizona, although it is entirely possible that dispersal, likely avian, was more gradational across the US, albeit never sampled. The placement of this New York and Arizona clade within a national context revealed that these strains represent the most ancestral extant strains of WNV in the United States.

The monophyletic nature of the nine Maricopa County NA/WN02 strains indicates that, at least for this lineage, there is most likely a mechanism that allows for viral overwintering in resident birds and/or *Culex* mosquitoes living in Maricopa County. Komar et al. [30] performed an extensive study in 2010 to identify Arizona-resident avian hosts of WNV. They found that communal roosting house sparrows (*Passer domesticus*), house finches (*Haemorhous mexicanus*), great-tailed grackles (*Quiscalus mexicanus*), and mourning doves (*Zenaida macroura*) account for the greatest proportion of resident bird infections. In addition to highly competent resident bird species in Maricopa County, both *Culex tarsalis* and the more abundant *Culex quinquefasciatus* vectors are present year-round in Maricopa County (Fig. 4). However, more extensive surveillance is needed during the winter season to better understand which mechanisms support overwintering. Although WNV has not historically displayed strong geographic clustering [31], the monophyletic clustering of the established Arizona strains is similar to that of those in Southern California [15]. Collectively, these studies indicate that the American southwest presents a suitable habitat for WNV to ecologically establish and persist across multiple years.

**Figure 4:**
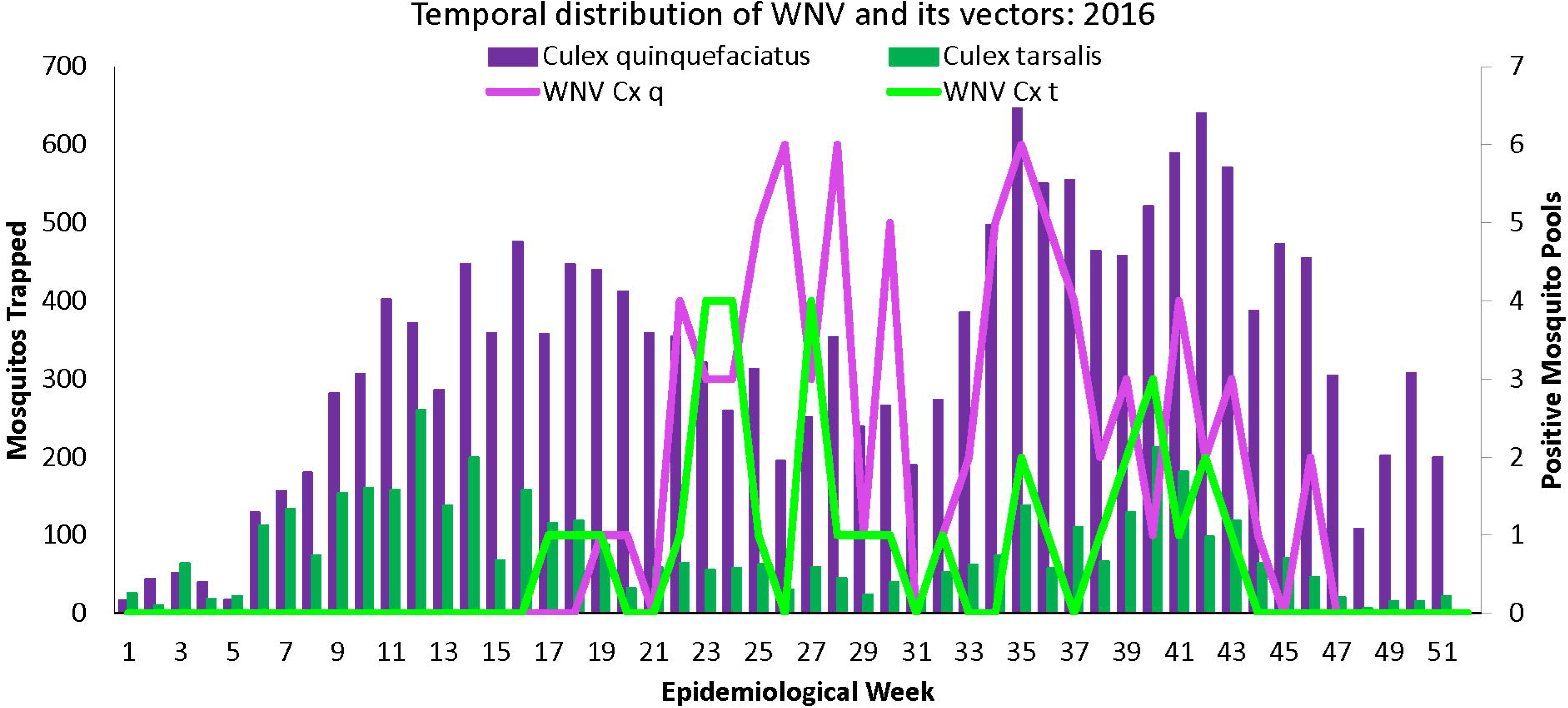
Distribution of *Culex* mosquitoes and WNV positive pools in Maricopa County by epidemiological week in 2016. The first y-axis represents the number of *Culex quinquefasciatus* (purple bars) and *Culex tarsalis* (green bars) individuals trapped each epidemiological week. The second y-axis represents the number of WNV positive *Culex quinquefasciatus* (purple line) and WNV positive *Culex tarsalis* (green line) mosquito pools trapped each epidemiological week.

While the two major lineages of WNV that circulate in the United States exhibit drastically different behavior with regards to establishment (NA/WN02) or transience (SW/WN03) in Arizona, their continued regional success is evident in sequence data from contemporary strains. Perhaps the most intriguing result of this study is that despite the diversity of Arizona strains over time, all carry the nonsynonymous mutation of NS5-K314R. This mutation appears to have arisen in 2001 (mean estimate: 2001.41, 95%CI: 2001.05-2002.18, Fig. S1), in the SW/WN03 lineage, and then independently arose on three separate occasions; twice in the NA/WN02 lineage and one additional time in the SW/WN03 lineage (Fig. S1). Given that previous selection analyses have identified this mutation as being positively selected [29, 32], we hypothesize that this mutation is important for adaptation to the Arizona ecosystem, and is at least beneficial to survival in other parts of the southwest United States.

---

**S1 Figure:** Tip-labelled maximum clade credibility phylogenetic tree reconstructed from 246 nationally distributed WNV genomes. Each tip consists of the accession number, vector or host where the strain was derived, two letter state code (or 4 letter code for location in Mexico), and the date of sampling to the nearest hundredth of a year. The gradient squares indicates the phylogenetic position of the NY99 (blue), NA/WN02 (red), and SW/WN03 (orange) genotypes. NA/WN02 and SW/WN03 strains are defined by the E-V195A mutation and the SW/WN03 genotype is defined by the NS4A-A85T mutation. Clade-defining mutations, as well as convergent mutations, are denoted by stars. Posterior probabilities are represented by the size (larger circles have higher values) and color (defined by the color legend) of circles at each node.

**Table Captions**

**Table S1:** Sequence metadata. The Genbank accession number, the host or vector mosquito that the strain was isolated from, the state of isolation, isolation date to the nearest hundredths of a year, major WNV clade the strain belongs to, clade-defining mutations, and notable convergent mutations are included. We adopted the host/vector naming scheme as in Pybus et al. [13]:**Hosts:** Ah, *Ardea herodias* (blue heron); Bj, *Buteo jamaicensis* (red-tailed hawk); Bu, *Bonasa umbellus* (ruffed grouse); Bvs, *Butorides virescens* (green heron); Cb, *Corvus brachyrhynchos* (common crow); Cc, *Cyanocitta cristata* (blue jay); Ccs, *Cardinalis cardinalis* (northern cardinal); Cl, *Columba livia* (pigeon); Dc, *Dumetella carolinensis* (catbird); Ec, *Equus caballus* (horse); Fa, *Fulica Americana* (American coot); Hs, *Homo sapiens* (humans); Pc, *Phoenicopterus chilensis* (Chilean flamingo); Ph, *Pica hudsonia* (black-billed magpie); Pn, *Pica nuttalli* (yellow-billed magpie); Px, *Phalacrocorax sp*. (cormorant); Qq, *Quiscalus quiscula* (common grackle); Zm, *Zenaida macroura* (mourning dove); **Vectors:** Cn, *Culex nigripalpus* (mosquito); Cp, *Culex pipiens* (mosquito); Cq, *Culex quinquefasciatus* (mosquito); Cs, *Culex stigmatosoma* (mosquito); Ct, *Culex torsolis* (mosquito); Cx, *Culex sp*. (mosquito). Dates were calculated using the collection dates’ corresponding day of the epoch calendar which was then divided by the total number of days in the year (365). The resulting decimal calculation was then added onto the collection year to the hundredths place.

**Table S2:** Tiled amplicon primer pairs. Primer pairs, the amplification pools to which they belong, primer lengths, melting temperature (Tm), GC content, and start and end position in relationship to the NY99 strain are included.

**Table S3:** Results from the path and stepping stone sampling analyses.

## Acknowledgements

Funding for this work was provided by the Arizona Technology Research and Initiative Fund (TRIF). Computational analyses were carried out using the Northern Arizona University High Performance Computing Cluster, Monsoon. We thank the Biodefense and Ecology Center and the Public Health and Clinical Translational Genomics Center, both of which are part of the Pathogen and Microbiome Institute, for helpful comments and suggestions during presentations of these results.

